# Development of Fully Human, Bispecific Antibodies that Effectively Block Omicron Variant Pseudovirus Infections

**DOI:** 10.1101/2023.03.07.531527

**Authors:** Jason Allen, Michelle Gonzalez, Jasbir Kaur, Melinda Smith, Junping You, Guojun Yang, Dongxing Zha, Ze Tian, Amin Al-Shami, Chunhua Shi, Jeffrey Molldrem, Tim Heffernan

**Author notes:** Contributions: AA, JA, CS, ZT, JM and TH wrote the manuscript. DZ, CS, JA, ZT, JM and TH designed the experiments. JA, MG, JK, MS, JY, GY, CS conducted the experiments and performed data analysis.

## Abstract

The emergence of highly immune invasive and transmissible variants of severe acute respiratory syndrome coronavirus 2 (SARS-CoV-2) has decreased the effectiveness of existing vaccines. It is, therefore, critical to develop effective and safe therapeutics for SARS-CoV-2 infections, especially for the most vulnerable and immunocompromised patients. Neutralizing antibodies have been shown to be successful at preventing severe disease from early SARS-CoV-2 strains, although their efficacy has diminished with the emergence of new variants. Here, we aim to develop fully human and broadly neutralizing monoclonal (mAb) and bispecific (BsAb) antibodies against SARS-CoV-2 and its variants. Specifically, we first identified two antibodies from human transgenic mice that bind to the receptor binding domain (RBD) of the SARS-CoV-2 spike protein and are capable of neutralizing SARS-CoV-2 and variants of concern with high to moderate affinity. Two non-competing clones with the highest affinity and functional blocking of ACE2 binding were then selected to be engineered into two BsAbs, which were then demonstrated to have relatively improved affinity, ACE2 blocking ability, and pseudovirus inhibition against several variants, including Omicron (B.1.1.529). Our findings provide one mAb candidate and two bsAb candidates for consideration of further clinical development and suggest that the bispecific format may be more effective than mAbs for SARS-CoV-2 treatment.

## Introduction

Since first identified in December 2019, Severe Acute Respiratory Syndrome Coronavirus 2 (SARS-CoV-2) has triggered the global Coronavirus Disease 2019 (COVID-19) pandemic that has, to-date, resulted in more than 5.3 million deaths. Like other coronavirus family members, SARS-CoV-2 is a single stranded RNA virus surrounded by a lipid envelop and four structural proteins [1]. Infection of the host cell is enabled primarily by the surface spike protein, which consists of homotrimers harboring two functional subunits: S1 and S2. Viral entry into the mammalian host cell is facilitated by the binding of the S1 subunit receptor-binding region (RBD) to the mammalian angiotensin-converting enzyme 2 (ACE-2), as well as by the cleavage of the spike protein by transmembrane protease serine 2 (TMPRSS2), which are both present on the mammalian host cell surface [4-5]. Accordingly, administration of a serine protease inhibitor capable of inhibiting TMPRSS2 [23] has been shown to block the entry of SARS-CoV-2 into host cells [5]. Both ACE-2 and TMPRSS2 are expressed on pulmonary, cardiac, renal, and intestinal tissue as well as the endothelium [2, 3], resulting in SARS-CoV-2 patients who present with concomitant respiratory and extrapulmonary symptoms. Further, and perhaps most concerning, SARS-Cov-2 has continued to accumulate mutations, especially within the S domain, that have resulted in the emergence of more infectious and deadly variants such as the SARS-CoV-2 Alpha (B.1.1.7), Beta (B.1.351), Kappa (B.1.617), Delta (B.1.617.2), as well as Omicron and its sub lineages [6]. To prioritize global monitoring and research, the World Health Organization (WHO) has classified emerging SARS-CoV-2 strains as variants of interest (VOI; Kappa and Mu variants) and variants of concern (VOC; Alpha, Beta, Gamma, Delta, and Omicron variants) (13, 14).

Efforts to develop effective therapies against SARS-Cov-2 remain ongoing. Initial attempts focused on designing therapeutic antibodies, with four monoclonal antibodies (mAb) receiving Emergency Use Authorizations (EUAs) from the United States Food and Drug Administration (FDA) for the treatment of SARS-Cov-2 [7] [1]. All four FDA-approved mAbs target the RBD and have been effective at treating mild-moderate SARS-CoV-2 infections and preventing progression to severe disease and/or hospitalization [7, 8], but have been shown to be relatively less effective against the highly contagious Omicron variant (24). The Omicron variant harbors at least 32 mutations in the spike protein, fifteen of which are in the RBD, as well as mutations in the membrane, envelope, and nucleocapsid proteins, which all contribute to the variant’s enhanced capacity to infect, multiply, and evade the immune response [9-11]. While initial efforts have shown that cocktails of two or more mAbs retain efficacy against different variants, this therapeutic approach is both cost- and time-intensive (25). As such, emerging efforts to neutralize SARS-CoV-2 variants have focused on engineering bispecific monoclonal antibodies (BsAb), which enable the simultaneous binding of two independent epitopes and have been used to treat a variety of pathologies, including cancer (26), autoimmune and inflammatory diseases (27), and viruses (28-29).

Here, we used humanized mice combined with single B cell cloning technology to generate and identify monoclonal antibodies that can bind and effectively neutralize the SARS-CoV-2 original virus and several of its mutants, including the Omicron variant. We also selected two of our antibody clones to design BsAbs that would recognize two different epitopes of the spike protein. Our results show a superior efficacy of the BsAbs in neutralizing the Omicron variant and preventing the infection of target cells.

## Materials and methods

### Covid-19 Immunization

Three ATX-GK+ female mice, 6-7 weeks old, were purchased from Alloy Therapeutics Inc. Blood was collected from each mouse. Each mouse was then immunized at the groin region with a 100 μl subcutaneous injection of a 1:1 (v: v) mix of RIBI adjuvant (S6322-1VL, Sigma) and 10 μg spike protein dissolved in PBS, and then boosted once per week for 3 weeks with the same regimen and injection sites. Antibody titers were measured right before the third boost. Three days after the third boost, we administered a final boost of 200 μl of the same mixture through an intraperitoneal injection. Mice were euthanized 4 days after the final boost. Blood, draining lymph nodes, spleen, and bone marrow were harvested. All animal experiments were approved by and conformed to the relevant regulatory standards of the Institutional Animal Care and Use Committee at MD Anderson Cancer Center (MDACC).

### Antibody discovery, top clinical lead selection, and BsAb construction

Alloy human transgenic mice were purchased from Alloy Therapeutics (**Waltham, MA**). Single B-cell Cloning (SBCC) technology was utilized to amplify variable heavy and variable light gene regions from immunized Alloy mouse memory B cells. The naturally paired variable regions were cloned into pcDNA3.1+ hIgG1 and hKappa expression vectors for subsequent ExpiCHO high-throughput transient expression in 96-well format. After Protein-A magnet beads purification, binding avidity was assayed using Bio-Layer Interferometry (BLI) with anti-Human IgG Fc capture (AHC) biosensors. Positive clones were then expressed at mg levels for further cell surface EC50, cross-reactivity, epitope binning, ligand blockage, and stability assays.

To generate bispecific 81×83 antibodies by fusion to heavy or light chains (Bs-A and Bs-B), heavy or light chain gene regions from clone 81 were cloned into hIgG1 or hKappa pcDNA3.1+ expression vectors respectively. Clone 83 single chain Fv domain was linked to either the clone 81 light chain (Bs-A) or heavy chain (Bs-B) N-terminus. Bispecific antibodies were expressed in ExpiHEK293 cells through transient transfection, and then purified using a Protein-A column (AKTA FPLC system) from ATUM (**Newark, California**). The binding of bsAb with different RBD mutations was also tested by BLI.

### High-throughput mAb production, purification, and analysis

High-throughput production of mAbs was performed by micro-scale transfection (1 mL) of paired heavy and light chain antibody sequences cloned into a single expression vector and expressed in Chinese hamster ovary (CHO) cells using the Gibco™ ExpiCHO™ Expression System and a protocol for deep 96-well plates (ThermoFisher Scientific). Briefly, synthesized antibody-encoding DNA (0.8 μg per transfection) was diluted in OptiPro serum free medium (OptiPro SFM), incubated with ExpiFectamine CHO Reagent, and added to 750 µL of ExpiCHO cell cultures into sterile 96 deep well plates using a ViaFlo 96 liquid handler (Integra Biosciences). Plates were placed in an Infors HT Multitron Pro incubator shaking at 1,000 rpm with 3 mm orbital diameter at 37°C in 8% CO2 and 80% humidity. The day after transfection, ExpiFectamine CHO Enhancer and ExpiCHO Feed reagents were added to the cells, followed by 6 days incubation at 32°C in 5% CO2 and 80% humidity.

Cells were harvested by centrifugation at 1500 x g for 10 min and supernatants transferred to new deep 96-well plates for high throughput micro-scale purification. Briefly, clarified culture supernatants were incubated with 15 µL MabSelect resin (Cytiva, formerly GE Healthcare Life Sciences) on a 3 mm orbital shaker at 1000 rpm at room temperature (RT) to capture mAb. The mixture was then transferred to pre-equilibrated fritted deep well filter plates, washed with PBS, eluted with 100 µL 50 mM phosphoric acid pH 3.0 into 96-well plates containing 15 µL neutralization buffer (20X PBS pH 11.0).

Recovered fractions were analyzed for yield, size, and purity using the LabChip® GXII HT Touch™ (Perkin Elmer, CLS138160) microfluidic CE-SDS platform and a ProteinEXact Reagent Kit (Perkin Elmer, CLS150466) according to the manufacturer’s protocol. Briefly, 2.5 μl of purified protein was mixed with 18 μl of sample buffer and 8.75 mM Iodoacetamide (Thermo Fisher Scientific, A39271) for non-reducing conditions. Samples were then heated to 70°C for 10 min, cooled to RT and mixed with 35 μl of water. Data analyses were then performed using GXII Reviewer software and calculations were based on the internal reference ladder.

### Mid-scale mAb production and purification

For relatively larger scale mAb expression, we performed transfections of CHO cell cultures using the Gibco™ ExpiCHO™ Expression System in 125 mL Erlenmeyer vented cap flasks (Corning) that contained 35 mL of ExpiCHO cells following the manufacturer’s protocol. Antibodies were purified from filtered culture supernatants by fast protein liquid chromatography (FPLC) on an ÄKTA Pure instrument using a 1 mL HiTrap MabSelect PrismA column (Cytiva, formerly GE Healthcare Life Sciences). Purified mAbs were concentrated and buffer changed into PBS using Amicon® Ultra-4 50KDa Centrifugal Filter Units (Millipore Sigma). Purified mAbs in PBS were then filtered using sterile 0.2-μm pore size filter devices (Millipore), and stored in aliquots at 4°C until future use.

### MAb quantification

To quantify purified mAbs, absorption at 280 nm (A280) was measured using a NanoDrop (ThermoFisher Scientific), and mAb concentration was calculated using the IgG sample type setting on the NanoDrop that uses a typical molar extinction coefficient for IgG.

### Detection of SARS-CoV-2-specific Ab titer in mouse serum

Wells of 96-well EIA high bind plates (Costar #3361) were coated with 2 µg/mL of SARS-CoV-2 S1 protein and His tag (Acro Biosystems #S1N-C52H4) at 4°C overnight. Plates were blocked with 1% non-fat dry milk and 1% BSA in PBS containing 0.05% Tween-20 (PBS-T) for 30 minutes at RT and then incubated for 1 hr at RT with serial dilutions of mouse serum in blocking buffer. After washing, bound antibody was detected using HRP-conjugated horse anti-mIgG (Cell Signaling #7076) for 1 hr at RT. After washing, plates were developed using TMB substrate (Thermo Fisher Scientific) and the reaction was stopped with H2SO4. The absorbance was measured at 450 nm using a CLARIOstar® Plus plate reader (BMG Labtech).

### Anti-SARS-COV2 mAb screening by BLI assay

BLI assays were performed on the Octet Red384 instrument at 30°C with shaking at 1,000 RPM. The mAb binding screen was performed after 96-well high-throughput expression and protein A purification. 100 nM of different SARS-COV2 clones were loaded to each of the AHC Biosensors for 600s. After loading, the baseline signal was recorded for 60 s in 10xkinetics buffer from the vendor. The sensors were then immersed in 10xkinetics buffer containing different concentrations of S1 (ACRO Biosystems), S1D614G (ACRO Biosystems), or RBD-mFc (ACRO Biosystems) recombinant proteins for 300 s. Dissociation was then measured for 300 s by immersing sensors in 10xkinetics buffer. As a control for non-specific binding, the background signal of empty sensors with loading BSA was subtracted at each time point.

### Kinetic analyses

For kinetic analyses for SARS-COV2-81, SARS-COV2-83, Bs-A, and Bs-B, antibodies were captured on AHC sensors; ligands were diluted to 100 nM in 10xkinestics and loaded for 600 s. After loading, the baseline signal was then recorded for 1 min in 10xkinetics. The sensors were immersed into wells containing S1-His or RBD-muFc with 100 and 20 nM in 10xkinetics buffer for 600 s (association phase), followed by immersion in 10xkinetics for an additional 600 s (dissociation phase). The background signal from each analyte-containing well was measured using empty reference sensors coated with same concentrations of BSA and subtracted from the signal obtained with each corresponding mAb loaded sensor. Kinetic analyses were performed at least twice with an independently prepared analyte dilution series. Curve fitting was performed using a 1:1 binding model and the ForteBio data analysis software. Mean k_on_ and k_off_ values were determined by averaging all binding curves that matched the theoretical fit with an R2 value of >=0.95. The germlines of each clone were calculated by IgBlast (https://www.ncbi.nlm.nih.gov/igblast/).

### Antibody competition binding assays and ACE2 blockage assays

RBD-MFc fusion recombinant protein (100nM) was captured onto AR2G biosensors (ForteBio) for 600 s. After loading and quenching of the active AR2G surface by 1 M ethanolamine, the baseline interference was then read for 60 s in 10xkinetics buffer. This was followed by immersion in a 100 nM solution of one SARS-COV2 antibody (mAB#1) for 600 s to saturate the binding signal, then 30 s for baseline in 10xkinetics, and then 300 s in 100 nM ACE2. The dissociation was then measured for 300 s by immersing sensors in 10xkinetics buffer. As a control for non-competing binding, mAb#1 was replaced as 10xkinetics. As a control for 100% competing, ACE2 was loaded the same as mAb#1. As a control for non-specific binding, the background signal of the binding of mAb to BSA-coated biosensors was subtracted at each time point.

### Pseudovirus assay

Lenti-SARS-CoV-2 Full Length Spike protein-pseudotyped (WT) and ACE2+ 293 cell lines were purchased from Genecopoeia. Viruses were produced as recommended by the manufacturer and harbored a luciferase expressing plasmid. Virus was concentrated using ultracentrifugation (95000 g for 3 hr) and suspended using OPTI-MEM (Lonza). Viral transfections assays were performed by plating target 293T (ACE2+ve) cells at 10000 cells/well in a white wall 96-well plate in DMEM supplemented with 10% FBS and allowing cells to adhere overnight at 37 °C. The next day, the virus was incubated with the indicated antibodies in complete media containing 8 µg/ml of polybrene for 1 hr on ice. Supernatants were removed from target 293 cells before adding the virus and antibody mix. Media was replaced the next day and levels of luciferase activity reflecting viral infection were measured at the 72 hr time point.

The pseudovirus assay for the Omicron variant was performed by Creative Diagnostic. Briefly, dilutions of the antibody sample at 3X the desired final concentration were prepared in cell culture media on 96-well plates. A virus control (VC) without the antibody sample was applied as the negative control. A SARS-CoV-2 positive control antibody (polyclonal, CD Cat# CABT-ST01) with neutralization response to both WT and Omicron (CD, B.1.1.529) was applied as the positive control. 50 µl of diluted antibodies and 50 µl of pseudovirus were incubated for 1 hr and then added to 4E05/well of Human ACE2 Stable Cell Line - HEK293T (CD Cat# CSC-ACE01) and incubated at 5% CO2 and 37°C for 48-72 hours. Luciferase activity, expressed in relative light units (RLUs), was determined according to the Luciferase assay system user’s manual.

## Results

Fully humanized mAbs were generated by immunizing ATX-GK+ female mice (Alloy Therapeutics) with the spike protein in PBS using SBCC technology (Supplemental Figure 1). A total of 152 clones with unique sequences were expressed, purified, and used in a BLI assay to characterize the S1, S1D614G, and RBD-mFc binding affinity of expressed mAb clones. In this BLI assay, the mAbs with a human Fc portion were initially captured by the anti-human IgG Fc (AHC) biosensors followed by an association reaction with the S1-His, S1D614G-His, or RBD-mFc analytes to assess the binding reaction and kinetics. Of the 152 final clones, 26 clones with unique sequences were identified based on positive binding to S1-His, the immunogen that had been used to vaccinate ATX-GK+ mice (Supplemental Table 1). Among these 26 clones, 14 clones were demonstrated to bind to S1D614G and the RBD-mFc, eight clones were shown to bind to S1D614G but not to the RBD-mFc, and four clones were shown to bind to S1 – likely around, but not on, the D614G position – but not to the RBD-mFc (Figure 1A). The 26 clones demonstrated a range of affinities to S1-His, with 17 clones exhibiting single digit nanomolar affinity, four clones exhibiting sub-nanomolar affinity, and five clones exhibiting double digit nanomolar affinities (Figure 1B). Epitope binning analysis of the RBD binding clones demonstrated that clones could be distributed into three different epitope bins. Specifically, binding to ACE2 could be blocked by clone #81, the sole occupant of Bin#2, but could not be blocked by clones from Bin #1 and Bin #3 (Figure 1C). Based on epitope specificities indicated by binning analysis findings and affinity data, clones #81 and #83, now respectively termed SARS-CoV-81 and SARS-CoV-83, were selected as the two final leads for the pseudovirus blockage assay. SARS-CoV-81 exhibited sub-nanomolar affinity while SARS-CoV-83 exhibited single digit nanomolar affinity (Figures 1D & 1E). Cell surface blockage assays demonstrated that SARS-CoV-81 could efficiently block RBD or S1-His binding to HEK293 cells that expressed human ACE2 (Figures 1F & 1G). Furthermore, comparison of SARS-CoV-81 with SARS-VHH-72, a single-domain antibody (VHHs) clone generated from a llama and capable of neutralizing SARS-CoV-2 [12], found that SARS-CoV-81 was more effective than VHH-72 at preventing ACE2 binding (Figures 1F & 1G).

**Figure 1.**
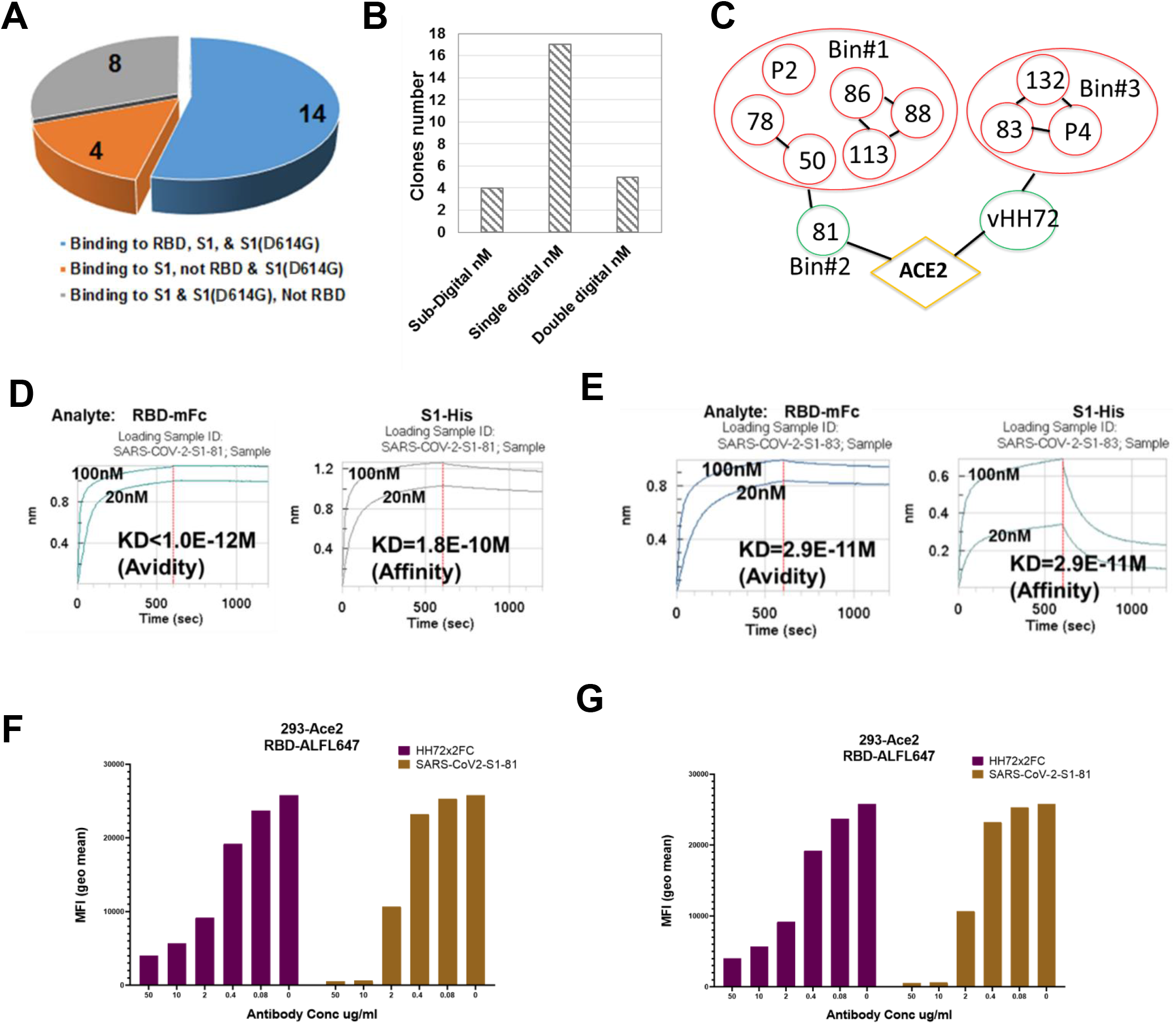
Development of anti-SARS-COV-2 antibody leads. (**A-B**) Distribution of screened clones that (A) bind to S1, S1D614G, and the receptor binding domain (RBD) and (B) exhibit affinity to S1-His using Bio-Layer Interferometry (BLI) analysis. (**C**) Binning map of representative clones binding to the RBD. (**D-E**) The avidity and affinity of (D) SARS-COV2-81 and (E) SARS-COV2-83 from the BLI assay. (**F-G**) Blockage of ACE2 binding on (F) HEK293 cells expressing S1 protein and (G) HEK293 cells expressing RBD by SARS-COV2-81 and vHH72.

Structural analyses of SARS-CoV-81 and -83 were then conducted using Molecular Operating Environment (MOE). Docking structure analyses found that the binding epitope of SARS-CoV-81 slightly overlapped with the ACE2 binding domain, and the binding epitope of SARS-CoV-83 did not overlap with the binding domains of either SARS-CoV-81 or ACE2 (Figure 2A). These data are consistent with our previous finding demonstrating that SARS-CoV-81 and SARS-CoV-83 do not compete when binding to the RBD or S1 and that SARS-CoV-81 blocks the binding of ACE2 to RBD and S1 (Figure 1). Receptor-binding domain residues L452, which has been shown to confer immune-escaping capabilities (22), as well as E484 and N501, which are destabilizing (22), were then mapped to the binding domains of clones SARS-CoV-81 and -83 as well as ACE2 (Figure 2B). Residue L452, E484, and N501 were found to be situated within the SARS-CoV-81 binding epitope (Figure 2C). Importantly, SARS-CoV-81 and -83 were shown to bind to different SARS-CoV-2 variants, including Beta (B.1.351), Gamma (P1), and Delta (B.1.617.2), without loss of affinity by BLI analyses (Fig 2D). However, loss of binding of SARS-CoV-81 to the L452R residue was observed (Figure 2D), with MOE modeling demonstrating a clear space overlap between the SARS-CoV-81 binding epitope and the RBD upon mutation of L452 to Arg (Figure 2C).

**Figure 2.**
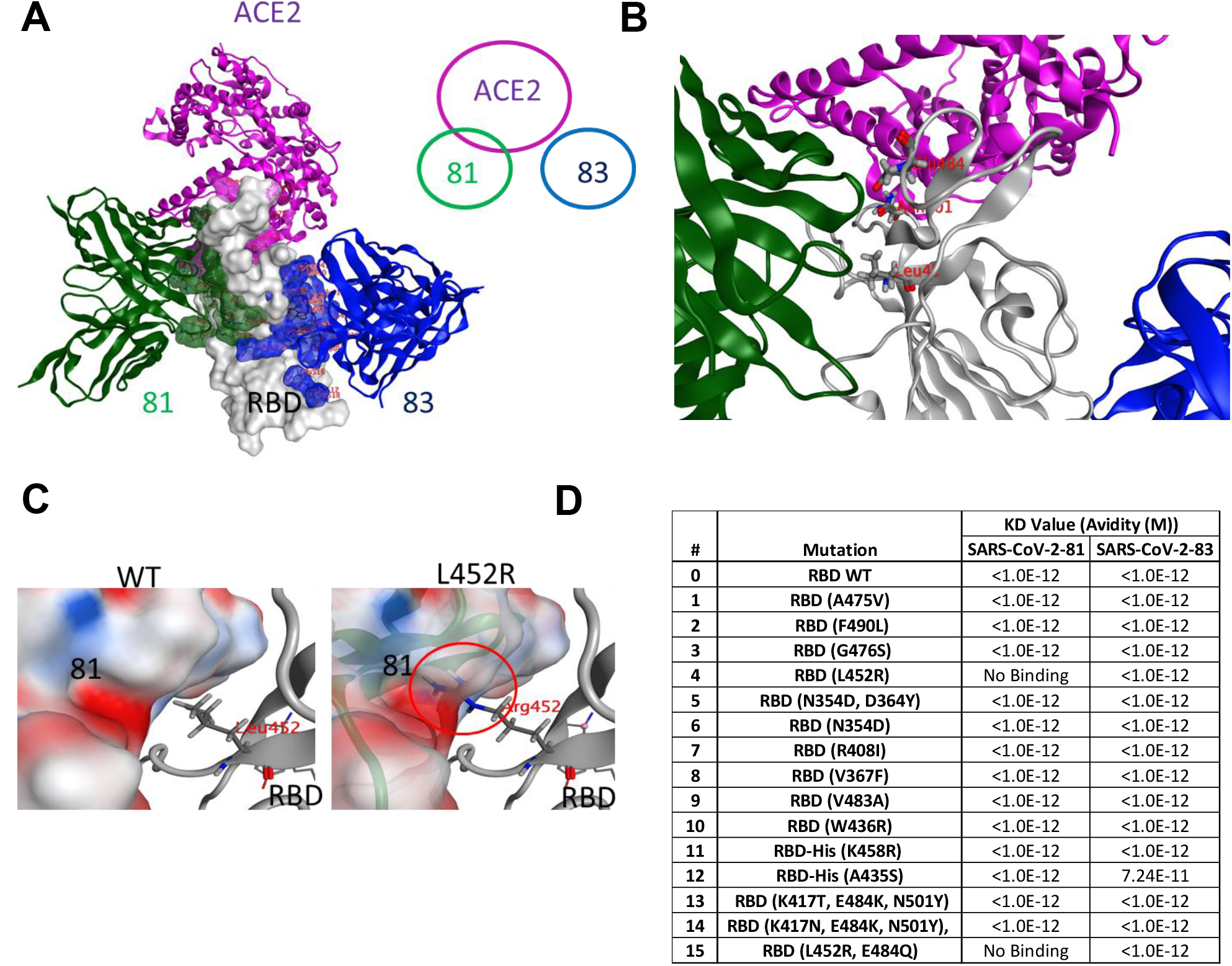
Docking of SARS-COV2-81 and SARS-COV2-81 receptor binding domain (RBD) binding epitopes to different RBD residues. **(A)** Docking structure of SARS-COV2-81 and SARS-COV2-83 by Molecular Operating Environment (MOE). The binding epitope of SARS-COV2-81, but not SARS-COV2-83, has a slight overlap with the ACE2 binding domain. **(B**) Magnification of the binding domains of SARS-COV2-81 and SARS-COV2-83 as well as the ACE2 binding domain. The L452, E484, and N501 residues, which are critical for enabling escape from antibody-based therapies, were mapped to the binding domains of SARS-COV-81 and SARS-COV-81. **(C)** SARS-COV2-81 cannot bind to a L452R-mutated RBD. *Left*, SARS-COV2-81 is capable of binding to wild-type (WT) RBD. *Right*, SARS-COV2-81 is not capable of binding to a RBD with the L452R residue. Red circle indicates a space between then SARS-COV2-81 RBD binding epitope and the L452R residue. **(D)** Binding affinity of SARS-COV2-81 and SARS-COV2-83 to RBDs harboring different mutations.

Due to their ability to recognize two non-overlapping epitopes, BsAbs have been hypothesized to enhance interaction with the target and may increase the efficiency of antibody-based anti-viral therapies. Here, we designed two BsAbs by adding the single chain Fv of SARS-CoV-83 to the light (Bs-A) or heavy (Bs-B) chains of SARS-CoV-81 (Figure 3A). Both BsAbs bound to the RBD of wild-type, Beta, Gamma, and Delta strains at picomolar avidity. Although SARS-CoV-81 did not bind to the L452R lineage, both Bs-A and Bs-B bound to all tested SARS-CoV-2 variants without losing affinity (Figure 3B). Further, both BsAbs were found to increase the blockage efficiency to the RBD of wild-type, Beta, Gamma, and Delta variants by a blockage assay (Figures 3C & 3D).

**Figure 3.**
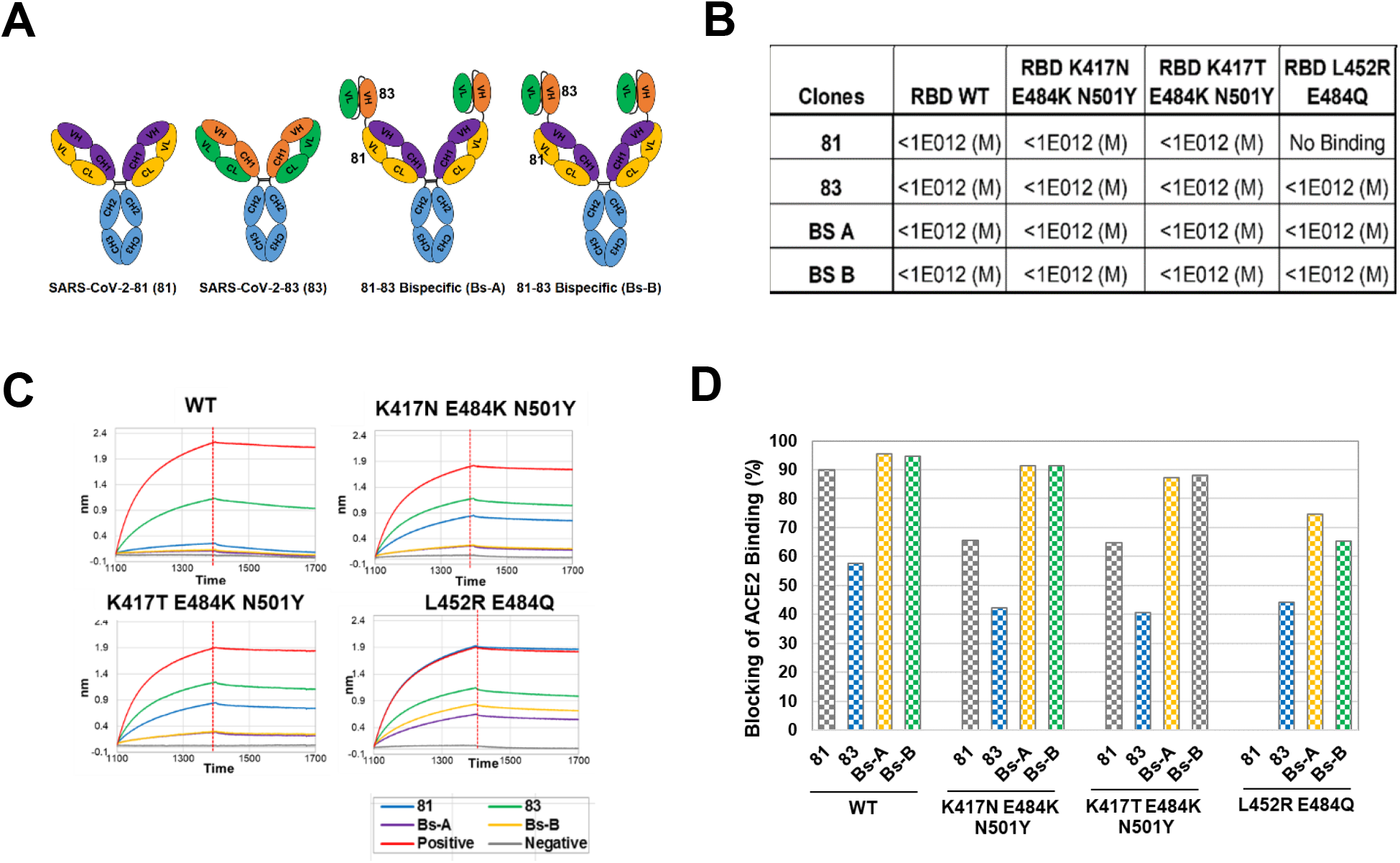
Bispecific antibodies (BsAb) increase the blockage of ACE2 binding to the receptor binding domain (RBD) of SARS-COV-2. **(A)** Schematic depicting the design of BsAbs, Bs-A and Bs-B. Briefly, the light (Bs-A) or heavy (Bs-B) chains of SARS-COV2-81 were added to the single chain Fv of SARS-COV2-83. **(B)** Characterization of SARS-CoV2-81, SARS-CoV2-83, Bs-A, and Bs-B binding to wild-type (WT) and mutated RBD of SARS-CoV2-2 by Bi-Layer Interferometry (BLI) analysis. **(C)** Capacity of Bs-A and Bs-B to block ACE2 binding to WT or mutant RBD of SARS-CoV2-2 by BLI assay. Positive control used is ACE2 binding to RBD directly without antibody. Negative control is 10xkinetics without ACE2 binding. **(D)** Comparison of the ability of SARS-CoV2-81, SARS-CoV2-83, Bs-A, and Bs-B to block ACE2 binding to WT or mutated RBD of SARS-CoV2-2.

To determine the ability of Bs-A and Bs-B to block spike-mediated infection of susceptible cells, we used a pseudovirus system consisting of a lentivirus with surface expression of the spike protein and a luciferase reporter. The resulting pseudovirus was capable of infecting 293 cells engineered to express the ACE2 receptor on the surface. In this system, the degree of viral infection can be measured by intracellular levels of luciferase activity. Escalating concentrations of the pseudovirus were incubated with 293 ACE2+ cells and SARS-CoV-81, SARS-CoV-83, BS-A, Bs-B, as well as with the negative isotype control Herceptin. Our findings show that Bs-A and Bs-B treatment offered significantly higher levels of protection against viral infection than that provided by mAb treatment, although SARS-CoV-81 offered partial protection (Figure 4). SARS-CoV-83, however, did not offer any protection (Figure 4).

**Figure 4.**
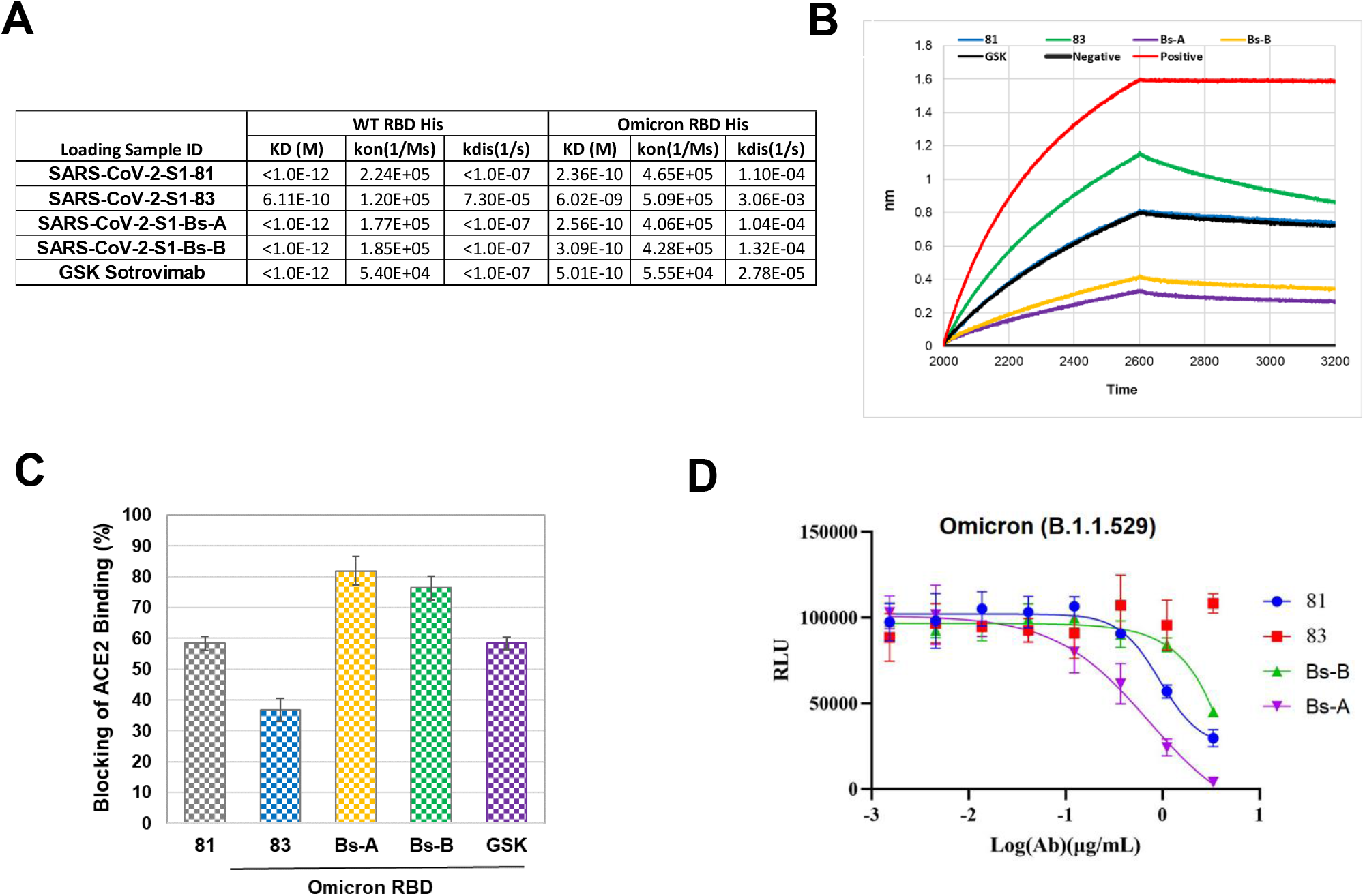
Capacity to neutralize the Omicron pseudovirus. **(A)** Binding avidity of SARS-CoV2-81, SARS-CoV2-83, Bs-A, Bs-B, and GSK Sotrovimab to the Omicron and wild-type (WT) receptor binding domain (RBD) recombinant protein. **(B)** Inhibition of recombinant ACE2 binding to the Omicron RBD by SARS-CoV2-81, SARS-CoV2-83, Bs-A, Bs-B, and GSK using Bi-Layer Interferometry (BLI) analysis. **(C)** Comparison of the ability of SARS-CoV2-81, SARS-CoV2-83, Bs-A, Bs-B, and GSK Sotrovimab to block ACE2 binding to the Omicron RBD using BLI analysis. **(D)** Characterization of the ability of SARS-CoV2-81, SARS-CoV2-83, Bs-A, and Bs-B to prevent Omicron pseudovirus infections in HEK293 cells expressing human ACE2.

We next assessed for the ability of Bs-A, Bs-B, SARS-CoV-81, and SARS-CoV-83 to neutralize the Omicron variant (B.1.1.529), which emerged as the dominant SARS-CoV-2 strain worldwide in late 2021 (14). Specifically, we measured the ability of our top lead mAbs and their bispecific formats to bind to the Omicron RBD, block ACE2 binding, and suppress Omicron pseudovirus infection. When compared to avidity with the RBD of wild-type SARS-CoV-2, SARS-CoV-81 and SARS-CoV-83 as well as their bispecific format bound to the Omicron RBD with a slight loss in avidity (Figure 4A). A similar loss of binding avidity was observed for GSK SotroviMab with the Omicron RBD (Figure 4A). Both Bs-A and Bs-B formats were observed to induce relatively high (∼80%) blockage of ACE2 binding, while SARS-CoV-81 and SARS-CoV-83 were found to induce moderate (∼60%) and low (∼35%), respectively, blockage of ACE2 binding by BLI analysis (Figures 4B & 4C). Of note, blockage efficiency of ACE2 binding was found to be similar between SARS-CoV-81 and GSK SotroviMab (Figures 4B & 4C). Findings from a pseudovirus blockage assay using HEK293 cells expressing human ACE2 demonstrated Bs-A to be capable of neutralizing the Omicron pseudovirus with a half-maximal inhibitory concentration (EC50) of ∼2.5 nM, while neutralization by BS-B and SARS-CoV-81 exhibited an EC_50_ of ∼160nM and ∼4.7nM, respectively (Figure 4D). SARS-CoV-83, however, was not able to neutralize the Omicron pseudovirus (Figure 4D). Overall, our findings indicate SARS-CoV-81 as either the standard IgG or bispecific format to be a promising candidate for further development as an antibody-based therapeutic strategy against Omicron and other SARS-CoV-2 strains.

## Discussion

As of April 2022, half a billion cases of SARS-CoV-2 have been confirmed worldwide, with more than 5.3 million deaths caused by the SARS-CoV-2 original strain and variants (13). Initial therapeutic development efforts resulted in the approval of EUA mAbs capable of neutralizing several SARS-CoV-2 strains, including VOCs (19) as well as prevent viral escape by targeting the RBD and obstructing engagement with ACE2 (20). However, SARS-CoV-2 mutates rapidly (20), with the RBD capable of accumulating many escape mutations while retaining the ability to bind to ACE2 (19, 21), thus limiting the efficacy of existing clinical mAbs and indicating a necessity for the rational design of non-competing mAb cocktails (20-21). Further, production of mAb cocktails is both time and cost-ineffective, with the possibility that the demand for anti-SARS-CoV-2 antibodies will outpace manufacturing capabilities acknowledged early in the COVID-19 pandemic (20).

Under the context of the rapid evolution of SARS-CoV-2 and the impractical manufacturing capabilities needed for developing antibody cocktails, recent efforts have focused on designing BsAbs to neutralize SARS-CoV-2 variants. The efficacy of BsAbs is thought to be more resilient than mAbs due to the hypothesis that BsAbs should retain efficacy even after the mutation of one binding site. Here, we first identified two fully human monoclonal antibodies, SARS-COV2-81 and SARS-COV2-83, capable of targeting the spike protein and RBD, and then developed our mAbs into two bispecific formats, Bs-A and Bs-B. Both SARS-COV2-81 and SARS-COV2-83 were demonstrated to be capable of binding with strong to moderate affinities to several SARS-CoV-2 variants, including Omicron and new variant lineages, but could not bind to the L452R mutant of the Delta (B.1.617.2) variant. Of note, SARS-COV2-81 exhibited avidity similar to GSK SotroviMab, a mAb approved as a therapeutic for patients infected with SARS-CoV-2 (30). Bispecific antibodies were then designed from combining the light (Bs-A) or heavy (Bs-B) chains of SARS-COV2-81 to the single chain Fv of SARS-COV2-83 and were demonstrated to bind to all tested SARS-CoV-2 variants, including the L452R lineage, with only a slight loss in avidity. Importantly, both Bs-A and Bs-B demonstrated superior avidity to Omicron RBD recombinant proteins when compared to that of SARS-COV2-81 or SARS-COV2-83 alone, and Bs-A was capable of neutralizing Omicron pseudovirus *in vitro*. Our findings align with other efforts to design BsAb against emerging and highly infectious SARS-CoV-2 variants. Specifically, when compared to parental mAbs, BsAbs have been shown to exhibit greater neutralizing activity against SARS-CoV-2 VOCS (15), including the Delta variant (17) and the Omicron variant and sublineages (16; 18), as well as have demonstrated a capacity to prevent escape mutants *in vitro* (15; 17-18). However, the rapid evolution of some SARS-CoV-2 variants has been shown to outpace the efficacy of available BsAbs (17), suggesting a need for the continual development of broadly neutralizing BsAb and/or BsAb that target epitopes different than those of existing BsAbs as potential treatments against emerging SARS-Cov-2 mutants.

In conclusion, we developed and characterized two fully human mAbs, SARS-COV2-81 and SARS-COV2-83, that were then used to design two BsAbs, Bs-A and Bs-B. Our data demonstrated enhanced neutralizing activity against VOCs, including Omicron and its sublineages, by Bs-B and Bs-B, but not SARS-COV2-81 and SARS-COV2-83. Of note, the avidity of our standard IgG SARS-COV2-81 to the Omicron RBD was comparable to that of GSK SotroviMab, thus suggesting SARS-COV2-81 to be a potential mAb candidate for further clinical development. Overall, our findings support the continual development and assessment of antibody therapy in the bispecific format as both efficacious and cost-effective therapeutic approaches for treating SARS-CoV-2 and its highly virulent variants.

## References

1. Yin, J., et al., Advances in the development of therapeutic strategies against COVID-19 and perspectives in the drug design for emerging SARS-CoV-2 variants. Comput Struct Biotechnol J, 2022. 20: p. 824–837.

2. Chavda, V.P., R. Pandya, and V. Apostolopoulos, DNA vaccines for SARS-CoV-2: toward third-generation vaccination era. Expert Rev Vaccines, 2021. 20(12): p. 1549–1560.

3. Hadizadeh, F., Supplementation with vitamin D in the COVID-19 pandemic? Nutr Rev, 2021. 79(2): p. 200–208.

4. Sagar, S., et al., Bromelain inhibits SARS-CoV-2 infection via targeting ACE-2, TMPRSS2, and spike protein. Clin Transl Med, 2021. 11(2): p. e281.

5. Hoffmann, M., et al., SARS-CoV-2 Cell Entry Depends on ACE2 and TMPRSS2 and Is Blocked by a Clinically Proven Protease Inhibitor. Cell, 2020. 181(2): p. 271–280 e8.

6. Mlcochova, P., et al., SARS-CoV-2 B.1.617.2 Delta variant replication and immune evasion. Nature, 2021. 599(7883): p. 114–119.

7. Anti-SARS-CoV-2 Monoclonal Antibodies. COVID-19 Treatment Guidelines. Available from: https://www.covid19treatmentguidelines.nih.gov/therapies/anti-sars-cov-2-antibody-products/anti-sars-cov-2-monoclonal-antibodies/.

8. Basu, D., V.P. Chavda, and A.A. Mehta, Therapeutics for COVID-19 and post COVID-19 complications: An update. Curr Res Pharmacol Drug Discov, 2022: p. 100086.

9. Dejnirattisai, W., et al., Omicron-B.1.1.529 leads to widespread escape from neutralizing antibody responses. bioRxiv, 2021.

10. Gupta, R., SARS-CoV-2 Omicron spike mediated immune escape and tropism shift. Res Sq 2022. rs.3.rs-1191837.

11. McCallum, M., et al., Structural basis of SARS-CoV-2 Omicron immune evasion and receptor engagement. Science, 2022. 375(6583): p. 864–868.

12. Wrapp, D., et al., Structural Basis for Potent Neutralization of Betacoronaviruses by Single-Domain Camelid Antibodies. Cell, 2020. 181(5): p. 1004–1015 e15.

13. Qianqian Li, M.Z., Ziteng Liang,Li Zhang,Xi Wu,Chaoying Yang,Yimeng An,Jincheng Tong,Shuo Liu,Tao Li,Qianqian Cui,Jianhui Nie,Jiajing Wu,Weijin Huang,Youchun Wang, Antigenicity comparison of SARS-CoV-2 Omicron sublineages with other variants contained multiple mutations in RBD. MedComm, 2022. 3(2).

14. World Helath Organization, WHO Coronavirus (COVID-19) Dashboard, https://covid19.who.int/.

15. De Gasparo R., et. al., Bispecific IgG neutralizes SARS-CoV-2 variants and prevents escape in mice. Nature. 2021 May;593(7859):424–428.

16. Wang., et. al., Biparatopic antibody BA7208/7125 effectively neutralizes SARS-CoV-2 variants including Omicron BA.1-BA.5, Cell Discov. 2023 Jan 7;9(1):3.

17. Li Z., et. al., An engineered bispecific human monoclonal antibody against SARS-CoV-2, Nature Immunology volume 23, pages 423–430 (2022).

18. Yuan M., et. al., A Bispecific Antibody Targeting RBD and S2 Potently Neutralizes SARS-CoV-2 Omicron and Other Variants of Concern, J Virol. 2022 Aug 24;96(16).

19. Dong J., et al., Genetic and structural basis for SARS-CoV-2 variant neutralization by a two-antibody cocktail, Nature Microbiology volume 6, pages 1233–1244 (2021).

20. Corti D., et al., Tackling COVID-19 with neutralizing monoclonal antibodies, Cell 184, June 10, 2021. 3086–3108.

21. Greaney A.J., et al., Complete Mapping of Mutations to the SARS-CoV-2 Spike Receptor-Binding Domain that Escape Antibody Recognition, Cell Host & Microbe, 2021 29, 44–57.

22. Fung K.M., et al., Antigen–Antibody Complex-Guided Exploration of the Hotspots Conferring the Immune-Escaping Ability of the SARS-CoV-2 RBD, Frontiers in Molecular Biosciences, 2022, 9, 797132.

23. Kawase M., et al., Simultaneous treatment of human bronchial epithelial cells with serine and cysteine protease inhibitors prevents severe acute respiratory syndrome coronavirus entry, J Virol. 2012 Jun;86(12):6537–45.

24. VanBlargan L.A., et al., An infectious SARS-CoV-2 B.1.1.529 Omicron virus escapes neutralization by therapeutic monoclonal antibodies. Nat Med. 2022 Mar;28(3):490–495.

25. Crowe Jr. J.E., Bispecific antiviral neutralizing antibodies are twice as nice. Nature Immunology volume 23, pages 346–347 (2022).

26. Rader C., Bispecific antibodies in cancer immunotherapy, Current Opinion in Biotechnology, 2020, 65, Pages 9–16.

27. Zhao Q., Bispecific Antibodies for Autoimmune and Inflammatory Diseases: Clinical Progress to Date, BioDrugs volume 34, pages 111–119 (2020).

28. Huang Y., et. al., Engineered Bispecific Antibodies with Exquisite HIV-1-Neutralizing Activity, Cell 2016, 165, 1621–1631.

29. Wec A.Z., A “Trojan horse” bispecific-antibody strategy for broad protection against ebolaviruses, SCIENCE, 354, 350–354.

30. FDA updates Sotrovimab emergency use authorization: https://www.fda.gov/drugs/drug-safety-and-availability/fda-updates-sotrovimab-emergency-use-authorization

