## Supplemental Data for "Development of Fully Human, Bispecific Antibodies that Effectively Block Omicron Variant Pseudovirus Infections"

*Corresponding Authors

**Supplemental Figure 1.** Timeline depicting the treatment schedule of humanized mice immunized at the groin region with a 100 μl subcutaneous injection of a 1:1 (v: v) mix of RIBI adjuvant (S6322-1VL, Sigma) and 10 μg spike protein dissolved in PBS. Briefly, blood was collected (prebleeding) from three ATX-GK+ female mice. Each mouse was then immunized and boosted once per week for 3 weeks. The final injection was administered 3 days after the last boost. Mice were then euthanized 4 days after the final boost, and blood, draining lymph nodes, spleen, and bone marrow were collected.

**Supplemental Table 1. Kinetics of all discovered clones from single B-cell cloning.**

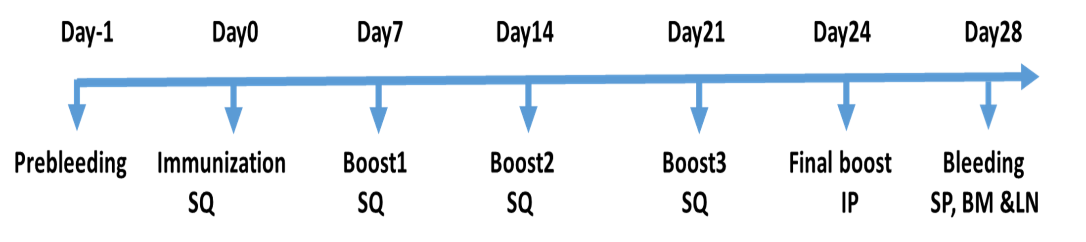
**Supplemental Fig.1 immunization and Abs generation summary**

**Supplemental Table 1. The avidity and affinity of 26 positive clones to SARS-CoV-2.**

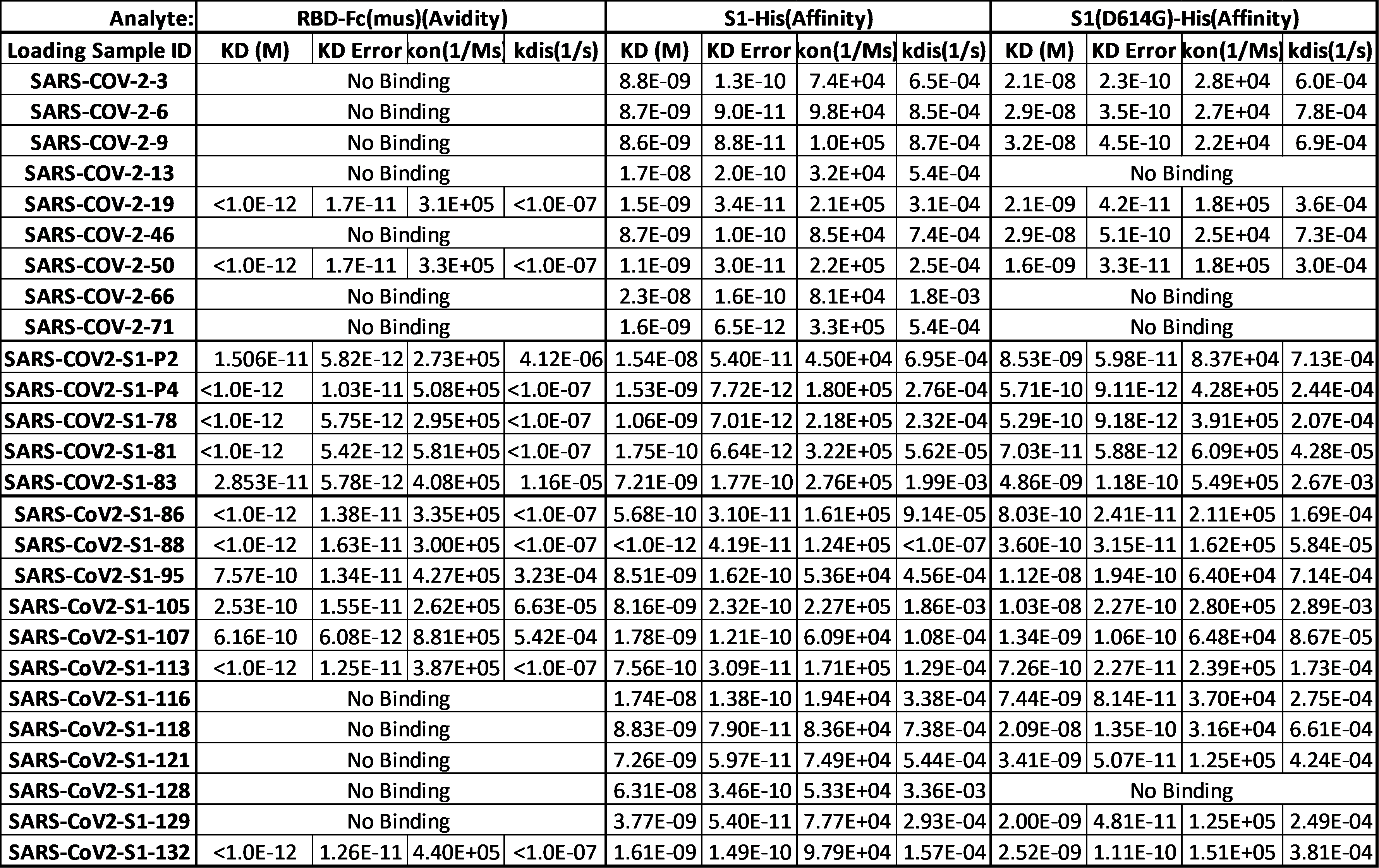
